# Tensing Flipper: Photosensitized manipulation of membrane tension, lipid phase separation and raft protein sorting in biological membranes

**DOI:** 10.1101/2024.06.25.599907

**Authors:** Joaquim Torra, Felix Campelo, Maria F. Garcia-Parajo

## Abstract

The lateral organization of proteins and lipids in the plasma membrane is fundamental to regulating a wide range of cellular processes. Compartmentalized ordered membrane domains enriched with specific lipids, often termed lipid rafts, have been shown to modulate the physicochemical and mechanical properties of membranes and to drive protein sorting. Novel methods and tools enabling the visualization, characterization and/or manipulation of membrane compartmentalization are crucial to link the properties of the membrane with cell functions. Flipper, a commercially-available fluorescent membrane tension probe, has become a reference tool for quantitative membrane tension studies in living cells. Here, we report on a so far unidentified property of Flipper, namely, its ability to photosensitize singlet oxygen (^1^O_2_) under blue light when embedded into lipid membranes. This in turn results in the production of lipid hydroperoxides that increase membrane tension and trigger phase separation. In biological membranes, the photo-induced segregated domains retain the sorting ability of intact phase-separated membranes, directing raft and non-raft proteins into ordered and disordered regions, respectively, in contrast to radical-based photo-oxidation reactions that disrupt raft protein partitioning. The dual tension reporting and photosensitizing abilities of Flipper enable simultaneous visualization and manipulation of the mechanical properties and lateral organization of membranes, providing a powerful tool to optically control lipid raft formation and to explore the interplay between membrane biophysics and cell function.

## INTRODUCTION

The plasma membrane is a complex assembly of lipids, proteins and sugars that separate the interior of the cell from the outside environment. Initially conceptualized as a fluid sea of lipids where proteins freely diffuse,^1^ it is now broadly established that specific components of the plasma membrane are spatiotemporally enriched and hierarchically organized in small domains of different lipid order and composition.^2–8^ Liquid ordered (Lo) regions, often termed as lipid or membrane rafts, are abundant in sphingolipids, cholesterol, and other lipids with saturated acyl chains, whereas liquid disordered (Ld) domains contain higher proportions of unsaturated lipids embedded in a more fluid environment.^9–12^ Such compartmentalization regulates the organization and interactions of a myriad of membrane protein receptors and is fundamental for a wide range of cellular processes such as signal transduction, trafficking, adhesion, motility, and many other cellular functions.^10,13–15^

The interplay between the formation and regulation of lipid domains, protein sorting and activation of cell processes has attracted much attention in the last years, fostering the development of methods and tools capable of visualizing or perturbing the organization of membranes. Smart fluorescent probes that sense changes in the biophysical and mechanical properties such as polarity or viscosity have become extremely powerful tools.^16,17^ However, other fundamental properties such as membrane tension, which plays a key role in the regulation of cell shape and motility,^18^ membrane fusion^19^, vesicle trafficking,^20^ among others,^21,22^ have remained particularly challenging to measure in living cells due to the lack of suitable fluorescent probes.^21^

In 2018, a fluorescent membrane tension reporter called Flipper-TR or Flipper was described^23^ and soon commercialized, becoming a reference tool in membrane tension studies in living cells.^24–29^ The mechanosensing ability of Flipper relies on the planarization of the two dithienothiophene (DTT) moieties (Figure S1a) in response to an increase in the lateral pressure.^23,30,31^ In non-confining environments, Flipper adopts a twisted conformation characterized by a broad absorption band centered around 400 - 450 nm and weak fluorescence emission. Insertion into lipid membranes sharpens and red-shifts the absorption band and activates the fluorescent properties, enhancing the emission intensity and increasing the lifetime (*τ*) as a function of the compression and planarization of the DTT scaffold.^32–35^ Particularly, Flipper *τ* is highly sensitive to its immediate environment, sensing subtle changes in membrane tension that can be detected and quantified using fluorescence lifetime imaging microscopy (FLIM).^23^

Despite its increasing popularity, previous works have also noticed manageable phototoxicity^30,36^ in living cells labeled with Flipper, although its causes and effects are not fully understood. Notably, no cell toxicity has been observed with Flipper incubated in dark conditions for more than 24 h,^23^ indicating that light is crucial for affecting cell viability. Light-induced cell damage is commonly associated with the formation of Reactive Oxygen Species (ROS) in a process called photosensitization, generally involving the triplet excited states of the fluorophores. Triplets are populated *via* intersystem crossing from the lowest singlet excited state and are more prone to react with nearby molecules and produce ROS favored by their longer lifespan.^37^ In turn, ROS rapidly react with lipids and proteins altering the normal functions of cells, causing severe cell damage when generated in essential organelles such as mitochondria or the plasma membrane.^38,39^ Of note, Flipper’s scaffold is rich in sulfur atoms, potentially enhancing the propensity of intersystem crossing due to the so-called heavy atom effect.^40^ Indeed, Flipper derivatives with high triplet yields have been reported recently.^30^ Therefore, it is highly plausible that Flipper excitation could lead to the production of ROS when embedded in membranes, although such photochemical process has not been reported so far.

The accumulation of Flipper in between the tails of the lipid that constitute cell membranes together with the high reactivity of acyl double bonds with ROS makes unsaturated lipids primary targets for ROS-mediated oxidation reactions. Photo-oxidation of lipids can proceed *via* electron transfer (Type I) or energy transfer (Type II) mechanisms yielding distinct oxidation products. Commonly, both mechanisms occur simultaneously in photosensitization processes, producing different oxidized lipid species and distinct effects on membranes as a function of the main ROS involved.^41^ In Type I, reactive radicals initiate a reaction cascade that can propagate far from the origin^42^ and lead to the formation of short-chain lipid aldehydes at double-bond sites.^43^ Experimental and computational studies have shown that truncated lipids reduce the viscosity of lipid bilayers, enhance membrane permeability and favor the formation of pores.^44–47^ In Type II processes, singlet oxygen (^1^O_2_) reacts with double bonds via the *ene* reaction producing lipid hydroperoxides,^48^ increasing the surface area, polarity and viscosity, while decreasing membrane thickness, elastic moduli and fluidity, yet preserving membrane integrity and impermeability.^43,46,47,49^

Lipid photo-oxidation also impacts the phase separation propensity in model and cell-derived membranes.^43,50^ Interestingly, different oxidized lipid species produced by distinct ROS in Type I or Type II mechanisms have been shown to alter the lateral organization of the membrane in opposite manners. While lipid hydroperoxides promote phase separation, truncated lipid aldehydes reduce lipid packing and favor lipid mixing.^45,46^ In this work, we aimed to assess whether Flipper indeed photosensitizes ROS when embedded in lipid membranes upon exposure to blue light and to unveil the potential effects on the properties and lateral organization in model lipid membranes as well as in biological membranes derived from cells.

## RESULTS

### Flipper embedded in lipid membranes generates ^1^O_2_under blue light inducing lipid hydroperoxidation

To investigate whether Flipper is capable of inducing photo-oxidation of unsaturated lipids via ROS production, we selected 1,2-dioleoyl-sn-glycero-3-phosphocholine (DOPC) and 1,2-dipalmitoyl-sn-glycero-3-phosphocholine (DPPC) as model examples of unsaturated and saturated lipids, respectively (Figure S1a), and prepared Giant Unilamellar Vesicles (GUVs). At room temperature, bilayers of DOPC are fluid and disordered whereas those composed of DPPC are gel-like and more ordered.^51^ As a result, Flipper exhibits red-shifted fluorescence excitation spectrum and longer *τ* when surrounded by DPPC as compared to DOPC (Figures 1a and S1b), consistent with previous data.^30,34^ Aqueous solutions of Flipper with the corresponding lipid were exposed to blue light (λ = 488 nm) for different times and we assessed the formation of lipid hydroperoxides using the Ferrous Oxidation–Xylenol orange (FOX) method.^52^ In this assay, hydroperoxides convert ferrous ions into ferric ions, forming a complex with xylenol orange that shifts its absorption from 430 nm to above 550 nm.^53^ Of note, the double bonds in the acyl tails in DOPC are prone to oxidation by ROS whereas DPPC is essentially inert.^41^ A new absorption band appeared at longer wavelengths in a light dose-dependent manner in DOPC samples. In contrast, the signal in DPPC remained unaltered (Figure 1b). The detection of hydroperoxides in unsaturated lipid solutions provided the first evidence for the formation of ROS upon Flipper photo-excitation.

**Figure 1.**
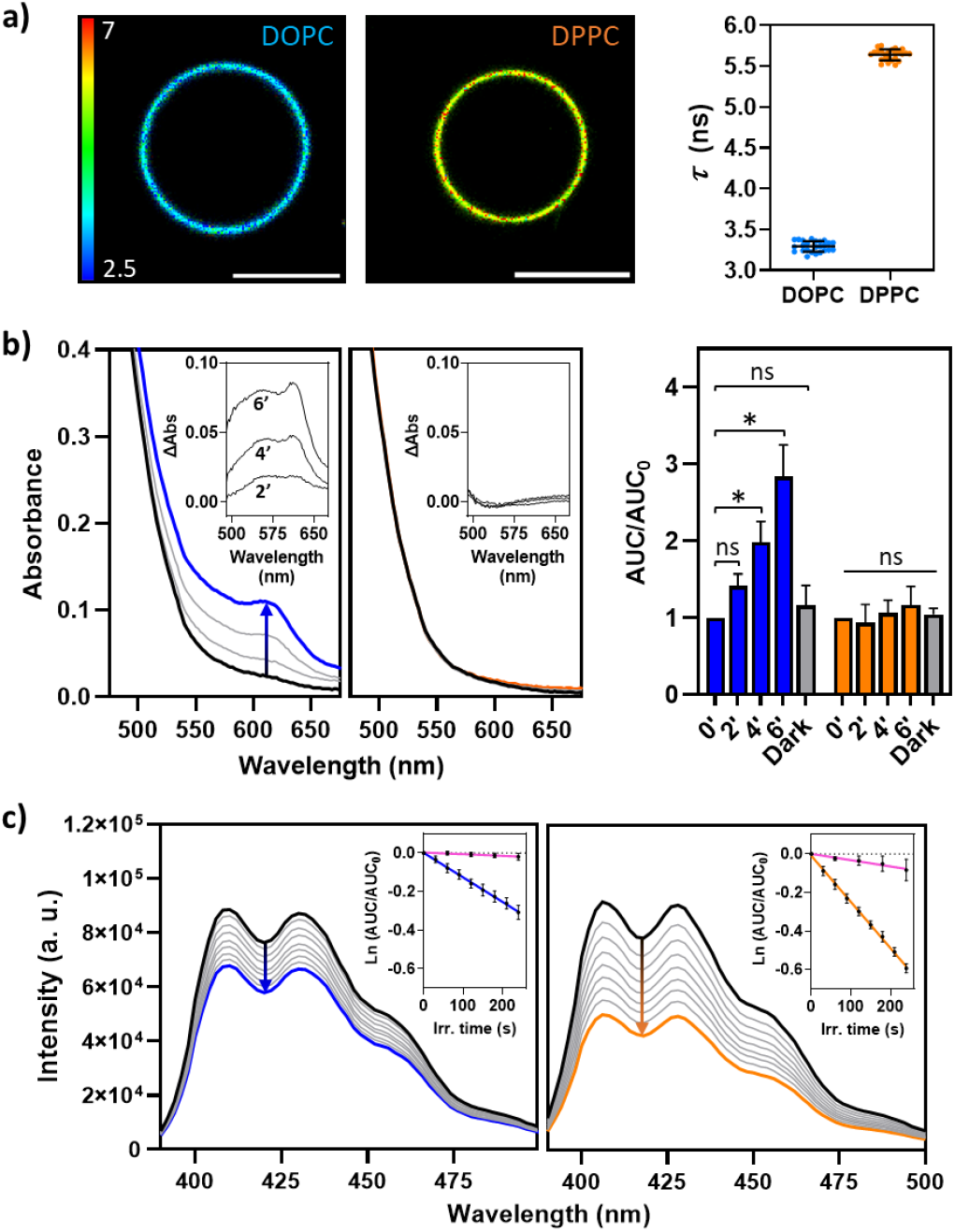
Mechanosensing and photosensitizing properties of Flipper. a) FLIM images of DOPC and DPPC GUVs labeled with Flipper (left) and the corresponding *τ* (right). Data from ≥ 25 GUVs from at least 3 independent experiments. Scale bars are 10 μm. b) Left: Changes in the absorption spectra of FOX incubated with solutions of 50 μg/ml DOPC (black to blue; first plot) or DPPC (black to orange; second plot) and 2.5 μM Flipper irradiated with blue light for different times. Solutions of Flipper and the corresponding lipid were incubated for 15 minutes in the dark prior to illumination. Insets show the difference spectra (ΔAbs) before and after 2, 4 and 6 minutes of irradiation. Right: Area under the FOX absorption curves (AUC) at different irradiation times (columns 2’, 4’ and 6’) integrated from 550 nm to 700 nm and normalized to non-illuminated solutions AUC_0_ (column 0). Gray columns dubbed dark indicate solutions of the corresponding lipid incubated with Flipper for 45 additional minutes (total 60 min) at room temperature in the dark. Data represent the mean ± SD of 3 replicates, \**P* < 0.01; ns: not significant determined by unpaired one-way ANOVA. c) Fluorescence photobleaching of 6 μM ADMA upon blue light irradiation of 2.5 μM Flipper in 50 μg/mL DOPC (left plot) or DPPC (right plot) D_2_O solutions. Insets: ADMA bleaching rates for DOPC (blue) and DPPC (orange), and in the presence of the specific ^1^O_2_ quencher NaN_3_ (10 mM, magenta). AUCs integrated from 390 nm to 500 nm at each irradiation time were normalized to AUC_0_, corresponding to non-illuminated solutions. Data are shown as mean ± SD of at least 3 replicates.

Lipid hydroperoxides are the primary products of photo-oxidation reactions of ^1^O_2_ with unsaturated lipids via Type II mechanism.^48,54^ Conversely, radical-based reactions in Type I mechanisms typically do not stop at the hydroperoxidation stage and further proceed producing truncated lipid aldehydes.^45^ To test whether Flipper photosensitizes ^1^O_2_under blue light, irradiation experiments were performed in DOPC and DPPC solutions supplemented with anthracene-9,10-diyl-bis-methylmalonate (ADMA), a fluorescent probe that selectively and irreversbly reacts with ^1^O_2_ yielding a non-fluorescent product.^55^ It is worth noting that ADMA is negatively charged and does not incorporate into the lipid bilayers but remains in the aqueous solution. Therefore, only the fraction of ^1^O_2_ that escapes from the hydrophobic core and reaches the aqueous media can be trapped by the probe. A common strategy to facilitate the detection of ^1^O_2_ is the use of deuterated solvents (i.e. D_2_O) to increase its lifetime in the aqueous solution.^56,57^ Blue light excitation of Flipper in bilayers of DOPC or DPPC in D_2_O induced a linear decrease in the emission intensity of ADMA in a light dose-dependent manner (Figure 1c). The bleaching of ADMA was more pronounced in DPPC membranes as compared to DOPC, which may reflect the consumption of ^1^O_2_ molecules upon reaction with the double bonds in DOPC, thus reducing the number of ^1^O_2_ molecules that can be trapped by ADMA. Moreover, we also note that Flipper absorbs a major fraction of the 488 nm light in DPPC due to the absorption red-shift induced by the more planar conformation adopted (Figure S1b). Finally, addition of sodium azide (NaN^3^), a well-established ^1^O_2_ scavenger,^58,59^ strongly reduced ADMA bleaching in both lipid solutions (Figure 1c). NaN_3_ is water soluble and thus outcompetes ADMA in quenching ^1^O_2_ molecules that escape from the lipid membranes and reach the aqueous media. Control experiments excluding either Flipper or light did not induce substantial changes in ADMA emission (Figure S1c). Taken together our results show that ^1^O_2_ plays a major role in the photo-oxidation of lipids induced by Flipper excitation.

### Lipid photo-oxidation induced by Flipper excitation increases the tension of unsaturated lipid membranes

The ability of Flipper to report on lipid membrane tension by means of its *τ* is widely exploited using FLIM.^30,36^ As a concentration-independent technique, FLIM is particularly well-suited to interrogating the close vicinity of the fluorescent probe and is broadly used to report changes in pH, concentration of ions and mechanical properties of membranes.^60^ To study the effects of Flipper excitation during FLIM imaging, GUVs made of DOPC or DPPC and labeled with Flipper were imaged in consecutive scans over more than 10 minutes. The excitation laser power was typically set to 25 % of the master power (*see methods*), which is comparable to that used in other Flipper-FLIM imaging experiments.^61^ Saturated DPPC GUVs were insensitive to Flipper excitation and remained essentially unaltered over the entire FLIM acquisition (Figures 2a, 2b and Video S1a). In strong contrast, DOPC GUVs exhibited notorious morphological changes within the first minutes of imaging, including an increase in the surface area accompanied by rapid fluctuations, formation of tubules and ultimately returning to a well-defined spherical shape of similar size as before illumination (Figure 2a and Video S1b). It is worth noting that we recorded consecutive single frames using 2x line accumulation in order to monitor progressive changes over long acquisition times. Notably, other works accumulated 10-25 imaging frames to obtain a FLIM image of Flipper,^36,61^ which, as observed in our experiments, may be already sufficient to alter the properties of the membrane of interest. The morphological changes observed are consistent with the effects of progressive formation and accumulation of lipid hydroperoxides produced by Type II mechanisms.^43^

**Figure 2.**
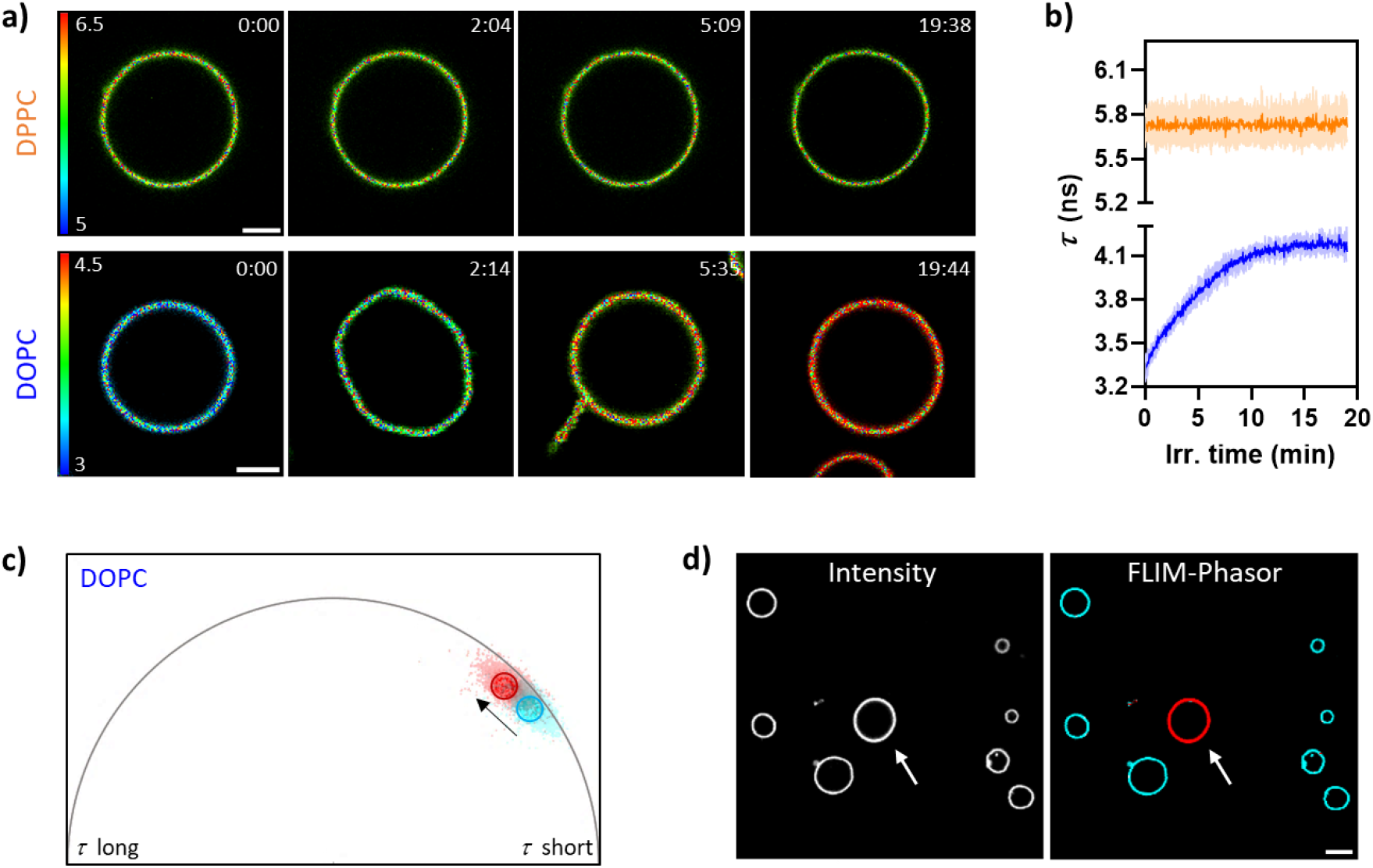
Photo-induced effects of Flipper excitation on model lipid membranes. a) Selected FLIM imaging frames over time of DPPC and DOPC GUVS labeled with 1 μM Flipper. Irradiation times are shown in min:sec on the top right of each frame. Scale bars are 5 μm. b) Time-course evolution of Flipper *τ* in DPPC (orange) and DOPC (blue) GUVs measured from the whole GUV at every imaging frame. Data are presented as mean ± SD for 5 independent GUVs. c) Phasor plot showing the shift in Flipper’s photon clouds in a DOPC GUV (from cyan to red) induced by prolonged FLIM imaging. d) Intensity (left) and FLIM-phasor images (right) showing a DOPC GUV imaged over multiple scans (white arrow) surrounded by others that were only exposed to a single imaging frame to capture the image, highlighting the highly localized effect of Flipper photosensitization at the illuminated region. Scale bar is 10 μm.

Strikingly, in DOPC GUVs, we also detected a gradual increase in Flipper *τ*, from 3.30 ns to around 4.15 ns where it reached a plateau (Figure 2b). Similar final *τ* values were obtained using different laser excitation powers ranging from 10 % to 75 % of the master power (Figure S2a) as well as in hydroperoxidized DOPC membranes produced using methylene blue and red light as described in Weber *et* al.^62^ (Figure S2b). These results indicate that the increase in *τ* originates from light-induced transformations of the lipids and not of the probe. The change in *τ* could also be monitored using the FLIM phasor plot analysis, producing a shift of the photon clouds towards longer *τ* (Figure 2c). Phasor plots are fit-free approaches that transform fluorescence decays from each pixel in a FLIM image to phasor coordinates, allowing to distinguish and separate different *τ* populations.^63^ We note that the photon clouds of Flipper fall inside the semicircle due to the bi-exponential nature of the decays.^23,36^ Moreover, the effects of Flipper excitation were highly localized and restricted to the illuminated region. This is consistent with the photosensitized production of ^1^O_2_ and its limited diffusion in aqueous media (<200 nm).^64^ As shown in Figure 2d, the increase in Flipper *τ* was only induced at the GUV exposed to multiple FLIM scans, leaving the surrounding lipid vesicles unaltered (Figure 2d). These results suggest that Flipper-induced lipid photo-oxidation could be attained with high spatiotemporal control.

Taken together, our results demonstrate that fluorescence imaging of unsaturated lipids with Flipper is sufficient to create oxidized species that progressively alter the composition and mechanical properties of the membranes. In addition, we show that hydroperoxidation of unsaturated lipids increases the tension of the lipid membrane as detected by the increase in Flipper *τ*. Moreover, such photo-induced effects can be monitored and characterized in real-time using the *τ* data. The combination of the photosensitizing and mechanosensing properties of Flipper therefore allows to simultaneously induce and visualize changes in the tension properties of the membranes.

### Simultaneous induction and visualization of lipid phase separation in model and biological membranes using Flipper

Previous reports have shown that lipid peroxidation favors phase separation in model lipid membranes, creating microdomains of different lipid order.^43,65–68^ Using model systems of ternary lipid mixtures incubated with compounds that produce ^1^O_2_ in high yield, segregated microdomains were formed *in situ* upon light exposure.^46,50,62^ Hence, we tested whether Flipper excitation is capable of inducing phase separation in synthetic lipid bilayers. We prepared GUVs assembled from well-characterized ternary mixtures of (1:1:1) 1-palmitoyl-2-oleoyl-sn-glycero-3-phosphocholine (POPC), DPPC and cholesterol (Chol), doped with saturated 1,2-Dipalmitoyl-sn-glycero-3-phosphoethanolamine labeled with far-red Atto-647N (DPPE-Atto647N) as a marker of Ld regions.^69,70^ Such lipid mixtures are known to form homogeneous membranes that phase separate upon hydroperoxidation at the single double bond present in the unsaturated POPC tail^46,50^ (Figure S1a). As a control, we also prepared and imaged GUVs composed of only POPC stained with Flipper, which showed homogeneous labeling with *τ* = 3.64 ± 0.03 ns (Figure S3a), slightly higher than in DOPC, which has two unsaturated acyl tails. Flipper excitation with blue light during extended FLIM imaging increased *τ* values, indicative of an increase in membrane tension (Figure S3b) and confirming that POPC is also susceptible to photo-oxidation by Flipper and blue light.

Dual-color confocal imaging of POPC/DPPC/Chol GUVs revealed a single phase in both DPPE-Atto647N and Flipper channels at initial illumination times (Figure 3, left column). Phasor plot analysis was used to group averaged lifetime (*τ*_ave_) components of Flipper for better differentiation of the photon clouds and visualization of the phases (Figure S3c). Continuous FLIM imaging triggered the formation of dynamic regions enriched with DPPE-Atto647N coinciding with a weaker fluorescence signal and shorter *τ*_ave_ from Flipper (Figure 3 middle and right columns and Video S2). These observations indicate lower membrane tension in Ld regions and are consistent with previous reports.^23,61^ Together, these results demonstrate that Flipper excitation progressively transforms the lipid composition of more complex membranes, eventually leading to phase separation. During the process, Flipper keeps sensing the properties of the bilayer and distinguishes Lo and Ld domains. Such dual inducer and reporter abilities using a single probe offer unprecedented potential to optically manipulate and characterize the mechanical properties of lipid membranes simultaneously.

**Figure 3.**
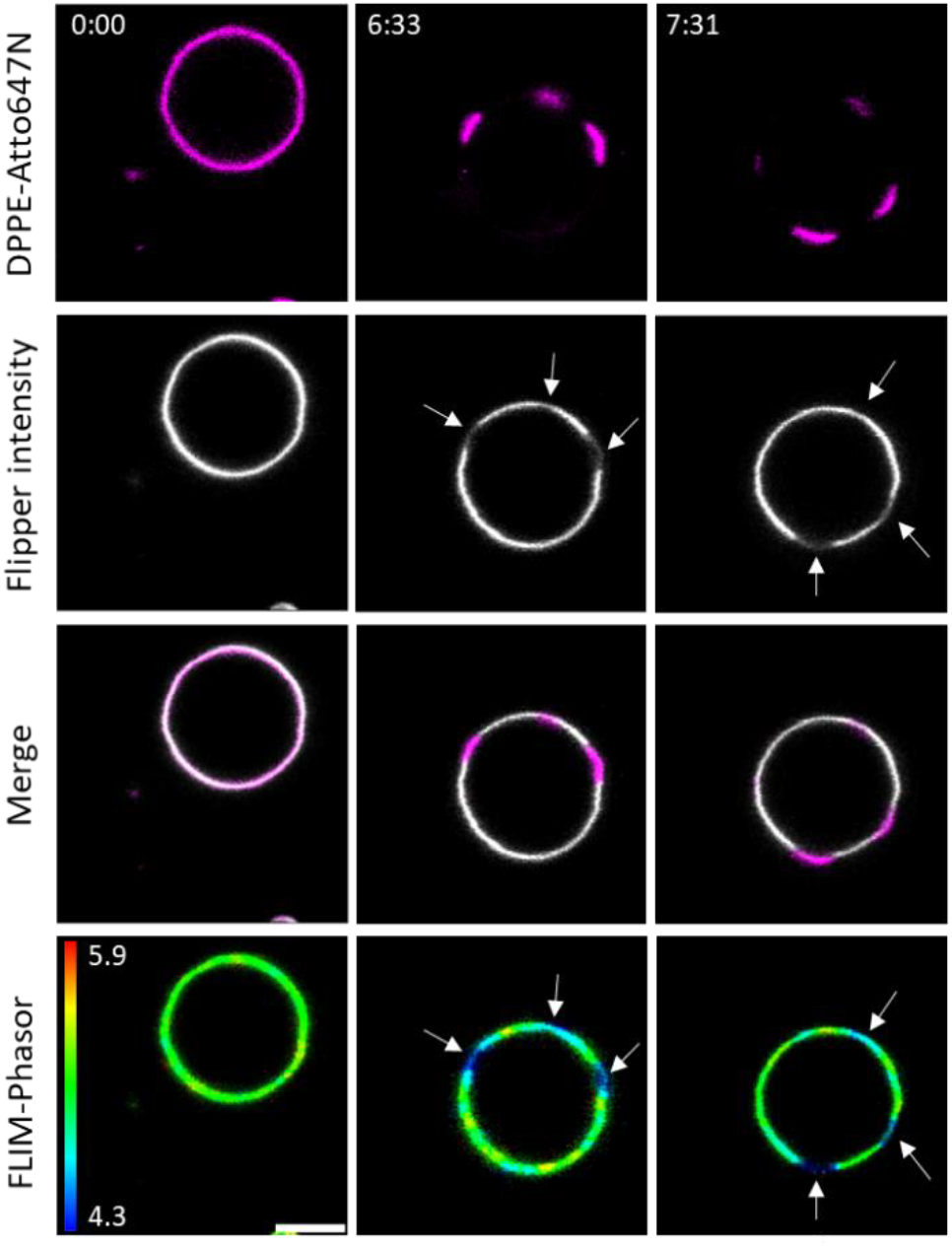
Flipper excitation triggers phase separation in model ternary lipid vesicles. Selected frames over time of a GUV composed of POPC/DPPC/Chol (1:1:1) and doped with the Ld marker DPPE-Atto647N transitioning from a single phase to segregated domains upon Flipper excitation in consecutive FLIM scans. Dynamic phases enriched with DPPE-Atto647N correlate with lower Flipper intensity and shorter *τ*_ave_ (pointed by white arrows), indicative of lower membrane tension in the Ld regions. Irradiation times in min:sec are shown on the top left. Scale bar is 5 μm.

While synthetic lipid bilayers are extremely useful membrane models offering strict control of the lipid composition, their biological relevance is inherently limited. Therefore, we next assessed the effects of Flipper excitation in Giant Plasma Membrane Vesicles (GPMVs) harvested from HeLa cells. GPMVs are attractive systems that closely mimic the compositional protein and lipid complexity of intact cell plasma membranes,^71–73^ which is particularly relevant in photo-oxidation studies considering the rich variety and abundance of unsaturated lipids at the plasma membrane. Moreover, the formation and the size of phase-separated domains can be tuned by the concentration of the vesiculating agents added during the preparation of GPMVs,^71,74^ enabling the isolation of non-phase-separated vesicles. Continuous FLIM imaging at 488 nm induced phase separation in GPMVs, which could be monitored in real-time by the differences in Flipper intensity and *τ*_ave_ (Figures 4a, S4a and Video S3). Brighter regions exhibited longer *τ*_ave_ whereas dimmer regions showed shorter *τ*_ave_, indicative of higher and lower membrane tension domains, respectively. Over time, the microdomains became larger and more defined, providing higher contrast between the phases in the *τ*_ave_ images.

**Figure 4.**
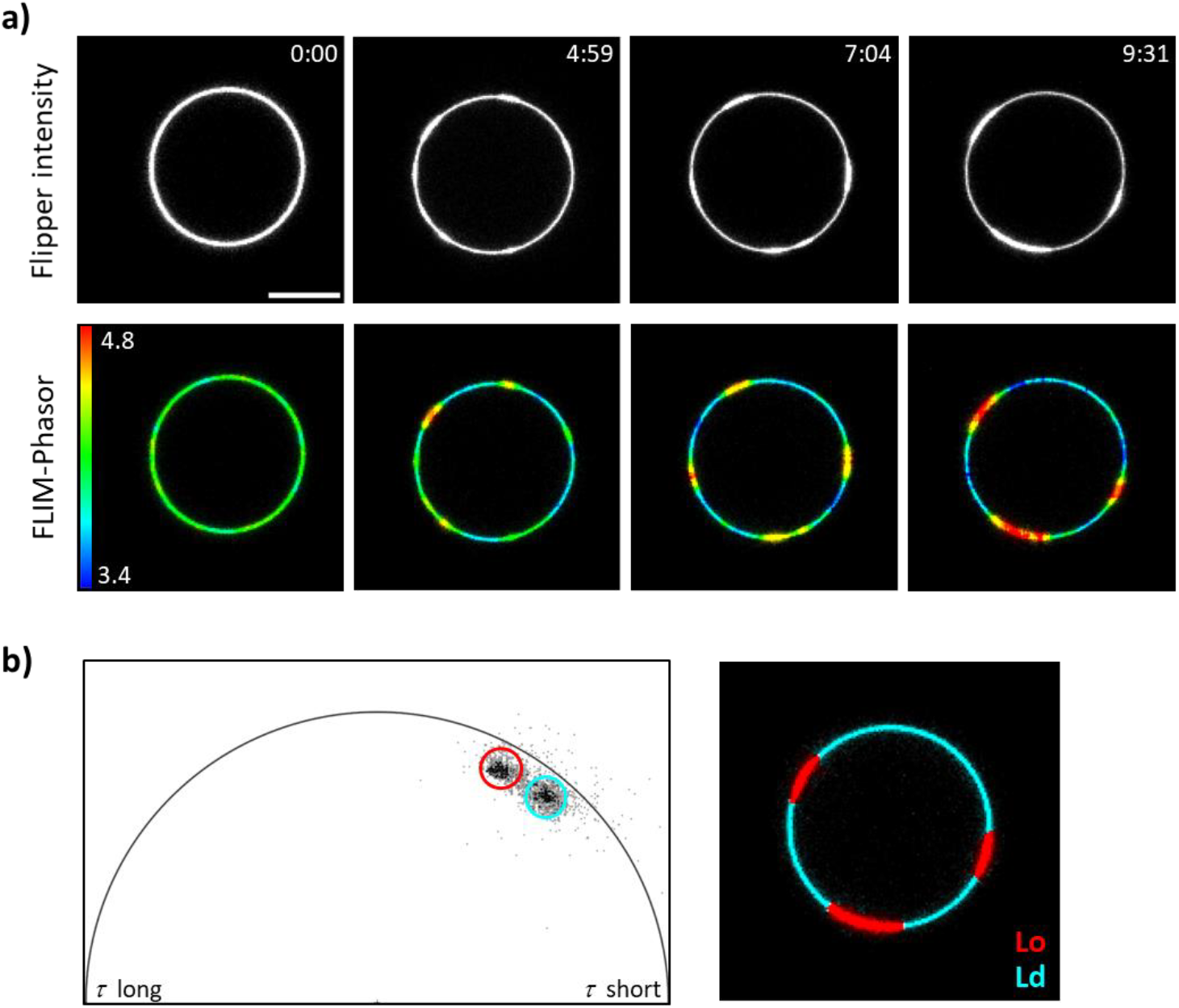
Photo-induced phase separation of biological membranes labeled with Flipper. a) Selected frames over time of a GPMV stained with 1 μM Flipper under prolonged FLIM imaging at 488 nm. The dynamic formation of small domains with different membrane tension can be observed in both intensity (up) and *τ*_ave_ (down) images. Irradiation times in min:sec are shown on the top right panels. Scale bar is 10 μm. b) Identification of Lo and Ld domains in photo-induced phase-separated GPMVs using the photon clouds in the Flipper phasor plot.

To unequivocally assign the response of Flipper with lipid order in GPMVs, we used the *bona fide* Ld marked DiD.^75,76^ Consistent with our results, DiD transitioned from homogeneous labeling to segregated accumulation in the low-tension regions upon photo-induced phase separation of the vesicles (Figure S4b). These results validate the ability of Flipper to distinguish between Lo and Ld phases without the use of additional markers (Figures 4b and S4c). Interestingly, an increase in Flipper *τ*_ave_ in the whole vesicle could be observed in most GPMVs prior to phase separation (Figures S5a and S5b). These results are fully consistent with the notion that membrane tension promotes phase separation in biological membranes as reported in previous studies.^77–80^ Altogether, our data show the ability of Flipper to photo-manipulate the mechanical properties and the lateral organization of biological membranes, while enabling the visualization and characterization of the newly segregated domains.

### Flipper-induced lipid photo-oxidation directs raft and non-raft protein segregation in biological membranes

The formation of segregated domains in lipid membranes drives the precise sorting and accumulation of membrane components into nanoscale domains or clusters that modulate a wide range of cellular functions.^8,14^ Indeed, a large number of membrane proteins have been identified to partition preferentially in either Lo or Ld regions, providing valuable information on protein behavior and in some cases, serving as established markers of raft and non-raft domains.^76^ For example, glycophosphatidylinositol-anchored proteins (GPI-APs) have been consistently found to be enriched in the Lo phase^11,81,82^ and are considered raft-markers, whereas several examples of trans-membrane proteins such as the transferrin receptor (TfR) are known to reside preferentially in Ld domains.^83,84^

Therefore, we next tested whether photo-induced phase separation of GPMVs into Lo and Ld domains using Flipper affects the organization of membrane proteins. Prior to GPMV formation, HeLa cells were transfected with Halo-tagged variants of GPI-AP or TfR (see *methods*) and labeled with the far-red dye Halo-JFX_650_. GPMVs containing the corresponding membrane protein were incubated with Flipper and a single frame was captured, showing homogeneous labeling in the two channels (Figure 5). Vesicles were then exposed to blue light from the excitation laser set at 25 % power until phase separation was induced (typically within 4-6 min of illumination) and single frames of the GPMVs were captured again in both channels. The fluorescence signal from GPI-APs and TfR showed segregation of the proteins, revealing that photo-induced formation of Lo and Ld domains directed the lateral partitioning of membrane proteins into specific regions. Notably, GPI-APs accumulated in Lo phases where Flipper exhibited higher fluorescence intensity and longer *τ*_ave_. In contrast, TfR was recruited into Ld domains characterized by low Flipper intensity and shorter *τ*_ave_. This behavior reproduces the preferential sorting of both proteins into raft and non-raft domains as observed in intact phase-separated membranes.^71,84^

**Figure 5.**
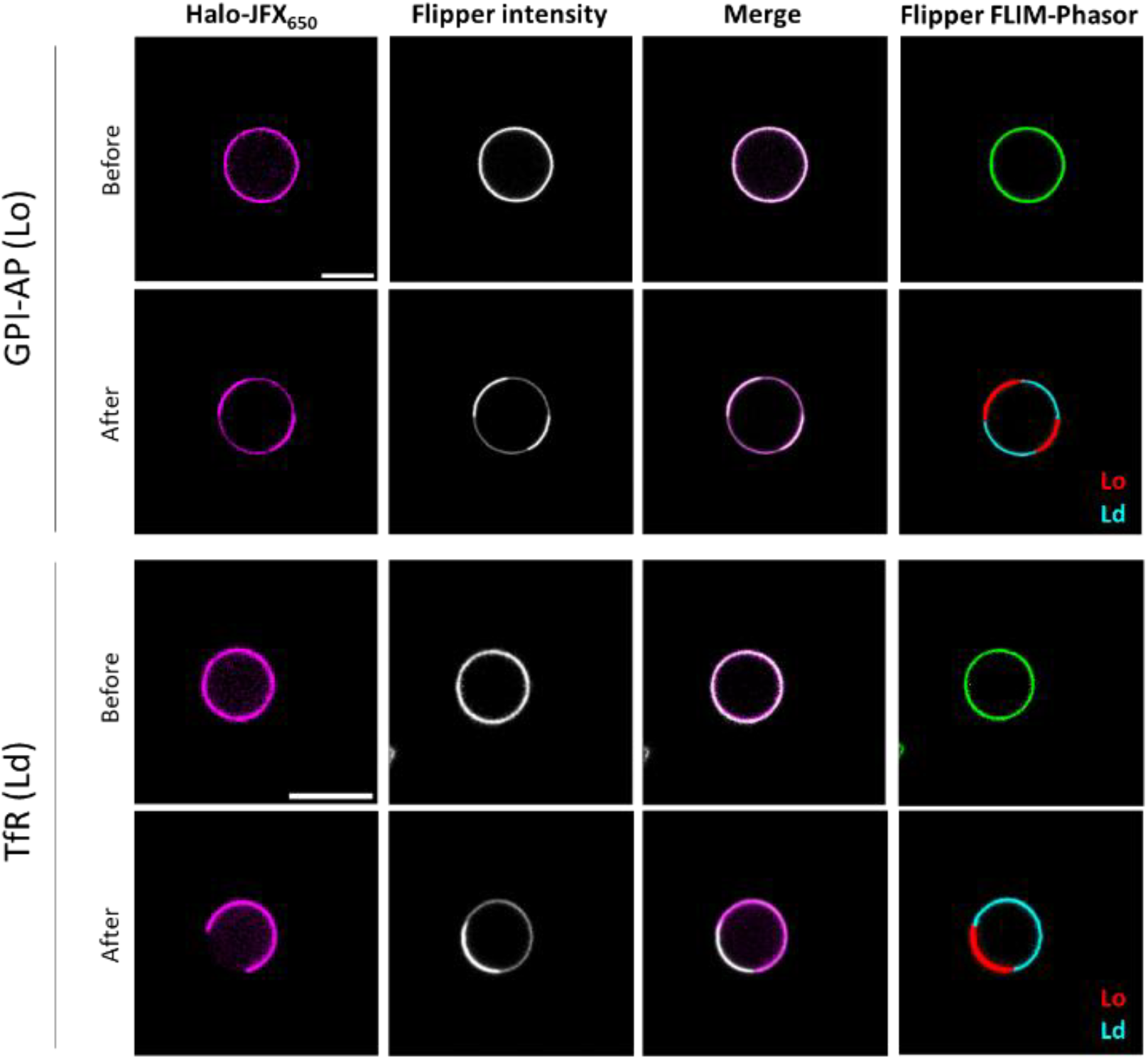
Photo-induced phase separation in biological membranes using Flipper directs raft and non-raft protein sorting into Lo and Ld domains, respectively. GPMVs expressing Halo-tagged variants of GPI-AP or TfR were labeled with 1 μM far-red Halo-JFX_650_ (magenta) and 1 μM Flipper (white). GPMVs initially showed a single phase in both channels (before) that separated after 4-6 min of blue light illumination (after). Scale bars are 10 μm.

Interestingly, a recent study found that radical-mediated lipid peroxidation *via* the Fenton reaction enhanced the phase separation propensity of GPMVs but disrupted the correct partitioning of raft proteins, which accumulated in Ld domains.^85^ Moreover, the authors showed that radicals produced oxidized lipids with truncated acyl chains, which decreased lipid packing in both Lo and Ld domains. The opposite behavior observed in our system therefore reinforces the notion that different oxidized lipid species divergently alter the properties and fate of the membrane, highlighting the role of the main ROS involved in distinct oxidation mechanisms. Altogether, these results demonstrate that illumination of Flipper embedded in biological membranes triggers the formation of co-existing domains of different lipid order, which retain the ability to direct the sorting of proteins into raft and non-raft membrane domains.

## DISCUSSION

It is well-recognized that membrane tension plays a crucial role in regulating essential cellular functions. The recent advent of mechanosensitive Flipper probes has provided a powerful tool for measuring membrane tension in model systems and in living cells, which has quickly boosted the use of Flipper in a broad number of studies. A few reports, however, have noticed manageable cell phototoxicity induced by Flipper under meaningful conditions,^30,36^ although its origin and potential consequences have not been further investigated. We rationalized that the sulfur atoms in the DTT scaffold could enhance the rate of intersystem crossing to the triplet excited state, favoring the production of cytotoxic ROS. Notably, unsaturated lipids are particularly prone to photo-oxidation reactions by ROS and low levels of oxidized lipids are sufficient to strongly alter the properties of the membranes at the nano- and microscopic scales^47,86^ before inducing obvious cell damage. In this work, we set out to assess the photosensitizing abilities of Flipper as well as its effects on model and biological membranes.

Using synthetic bilayers of DOPC or DPPC as selected examples of unsaturated and saturated lipids, respectively, we found that Flipper produces ^1^O_2_ when exposed to blue light. Moreover, we also detected lipid hydroperoxides only in illuminated DOPC membranes incubated with Flipper. Hydroperoxides produced at the double bond sites of the lipid acyl chains occupy a larger volume and tend to migrate to the aqueous interface due to their hydrophilic character, causing membrane fluctuations and increasing the surface area.^87^ Buds and tubules are also formed to accommodate the excess area and the vesicles ultimately recover the spherical size. On the microscopic scale, lipid hydroperoxidation increases membrane viscosity and polarity, decreases membrane thickness, elastic moduli and fluidity, while membrane permeability is retained.^41,47^ Conversely, truncated lipid aldehydes produced *via* Type I oxidation cause a large decrease in membrane viscosity, while increase membrane permeability and the propensity of pore formation.^44^

FLIM is the fluorescence microscopy technique of choice to study membrane tension as a function of Flipper *τ* but it commonly requires high and/or prolonged excitation conditions to collect a sufficient number of photons for accurate *τ* determination. We evaluated the effects of multiple FLIM imaging scans at 488 nm on model DPPC and DOPC GUVs labeled with Flipper. We found that DPPC was insensitive to photo-oxidation and remained es-entially unaltered. In contrast, DOPC GUVs exhibited notorious morphological changes in excellent agreement with the reported effects induced by ^1^O_2_-based lipid hydroperoxidation.^43^ Moreover, Flipper *τ* increased gradually throughout FLIM imaging, indicative of an increase in the tension of the oxidized membranes and providing direct evidence to link ^1^O_2_-mediated lipid hydroperoxidation with an increase in membrane tension.

Using ternary mixtures of lipids that form microscopically homogeneous membranes, phase separation was induced by prolonged Flipper excitation, forming coexisting domains of different lipid order as revealed by an Ld marker. Notably, Flipper also distinguished membrane phases, exhibiting higher fluorescence intensity and longer *τ* in Lo regions, in agreement with reported data.^23,61^ These results further demonstrate that Flipper is able to gradually modify the composition of lipid membranes under blue light, causing the reorganization of the different components and eventually leading to the formation of segregated regions.

Lipid phase separation dynamically drives the lateral organization of cell membranes, forming compartmentalized regions that regulate cell functions.^88^ Of note, the rich abundance and diversity of unsaturated lipids in cell membranes make them particularly prone to photo-oxidation reactions. To study the effects of Flipper in biological membranes, we used GPMVs derived from HeLa cells that retain the complex lipid and protein composition of the plasma membrane. GPMVs showed homogenous staining with Flipper at the beginning of FLIM imaging but phase-separated upon prolonged exposure to blue light. Notably, Flipper unambiguously distinguished Lo and Ld domains.

Photo-induced lipid segregation has been previously observed in artificial membranes and explored as a method to mimic lipid raft formation and to assess their biological implications.^43,89^ In a very recent study using GPMVs, lipid peroxidation induced chemically by highly reactive radicals (Fenton reaction) enhanced the phase separation propensity of GPMVs.^85^ The peroxidation process led to the formation of truncated lipid aldehydes that accumulated preferentially in the disordered phase but also decreased lipid packing in both Lo and Ld domains.^85^ Notably, peroxidized GPMVs lost the ability to segregate raft proteins into Lo regions, which accumulated in disordered phases. On the other hand, non-raft proteins remained in the Ld phase.^85^ Similarly, a previous study reported the disruption of GPI-APs nanoplatforms at the plasma membrane upon the addition of truncated oxidized phospholipids.^90^ In contrast, here we observed that the microdomains formed upon Flipper excitation retained the protein sorting ability, directing raft and non-raft proteins to the Lo and Ld regions, respectively. The opposite behavior reasonably arises from the intrinsic reactivity of the main ROS involved in distinct lipid oxidation mechanisms, leading to the formation of different oxidized lipid species. As described above, the individual contribution of truncated aldehydes or hydroperoxides has paramount importance on the properties of lipid membranes and raft homeostasis, inducing in most cases antagonistic effects.^46,50^ In future studies, lipid photo-oxidation methods with predominant Type I or Type II character can be further explored and even combined to tune the properties of biological lipid membranes and manipulate raft homeostasis. Particularly, photosensitized reactions are advantageous providing high spatiotemporal control on the formation of ROS and the oxidative effect, in contrast to the bulk production and indiscriminate activity induced by chemical methods.

The biological consequences as well as potential applications of Flipper photosensitization remain to be fully determined. Since membrane tension governs fundamental cell functions, subtle changes perturbing membrane homeostasis may strongly impact the lateral organization of cell-surface proteins and alter downstream signaling. Indeed, tools capable of precisely tuning the distribution of membrane proteins are in high demand.^88,91–93^ The dual tension reporting and photosensitizing abilities of Flipper may be considered in light-based tension measurements since the effects can be notorious within a few imaging frames captured under standard FLIM acquisition conditions.^36,61^ In fact, caution should be taken when performing prolonged fluorescence imaging of lipid membranes in air-saturated solutions (i.e., in the presence of molecular oxygen). Steady photosensitization of ROS by conventional and newly developed smart membrane probes over time, even in very low yields, will progressively alter the lipid environment and thus compromise the fluorescence readout. However, such abilities also offer great potential for dynamically visualizing and challenging membrane heterogeneity with high spatiotemporal control. All in all, the development of improved systems coupling photosensitized lipid oxidation with sensing membrane properties may provide powerful tools to study the interplay between the lateral organization and biophysical properties of the membrane with cell functions.

## CONCLUSION

The uses of Flipper, a commercially available fluorescent reporter of membrane tension using FLIM imaging are steadily growing. In this work, we showed that Flipper produces ^1^O_2_ when embedded into lipid membranes and exposed to blue light, inducing the hydroperoxidation of unsaturated lipids. This process causes notorious morphological changes, an increase in membrane tension and triggers phase separation in model and biological membranes. The photo-induced microdomains in cell-derived membranes exhibit different lipid order and direct raft and non-raft membrane proteins into Lo and Ld phases, respectively. These findings highlight the different behavior of distinct oxidized lipid species and emphasize the potential of photosensitized lipid peroxidation for precise manipulation of membrane lateral organization and exploring how membrane biophysics regulate cell function. Our results uncover additional properties of Flipper that may impact light-based tension measurements and open new avenues for optically controlling and visualizing lipid raft formation and directed membrane protein sorting.

## Author Contributions

The manuscript was written through contributions of all authors. All authors have given approval to the final version of the manuscript.

## Notes

The authors declare no competing financial interest.

## ACKNOWLEDGMENT

The authors thank Prof. Jennifer Lippincott-Schwartz for kindly providing the Halo-tagged GPI-AP and TfR RUSH plasmids (Addgene plasmids #166903 and #166905, respectively), Merche Rivas for cell culture support and Maria Marsal for fluorescence imaging assistance.

