## Supporting Information for "Tensing Flipper: Photosensitized manipulation of membrane tension, lipid phase separation and raft protein sorting in biological membranes"

### Materials and methods.

#### Materials:

Lipid solutions in $CHCl_3$ of 1,2-dioleoyl-sn-glycero-3-phosphocholine (DOPC), 1,2-dipalmitoyl-sn-glycero-3-phosphocholine (DPPC), 1-palmitoyl-2-oleoyl-sn-glycero-3-phosphocholine (POPC), 1,2-dipalmitoyl-sn-glycero-3-phosphoethanolamine-N-(cap biotinyl) were purchased from Avanti Polar Lipids. Atto647N-labeled 1,2-Dipalmitoyl-sn-glycero-3-phosphoethanolamine (DPPE-Atto647N) was obtained from ATTO-TEC, Cholesterol (Chol) ≥ 99% from Sigma-Aldrich, Neutravidin from ThermoFisher Scientific.

Flipper-TR® (Flipper) was purchased from Spirochrome, $DiIC_{18}(5)$ solid (DiD) from Molecular Probes, Life Technologies Corporation, and Halo-tagged $JFX_{650}$ from Promega. Deuterated water ($D_2O$), 9,10-Anthracenediyl-bis(methylene)dimalonic acid (ADMA), sodium azide ($NaN_3$), polyvinyl alcohol (PVA, mw = 145 kDa), paraformaldehyde (PFA) and dithiothreitol were obtained from Sigma-Aldrich. Ferrous ammonium sulfate hexahydrate >99% and xylenol orange (XO) sodium salt were purchased from Fischer Scientific.

#### Ferrous Oxidation–Xylenol orange (FOX) assay for detection of lipid hydroperoxides

Flipper (2.5 µM) was added to lipid solutions of 50 µg/mL DOPC or DPPC in $H_2O$, incubated in the dark for 15 min and then exposed to 488 nm light illumination for 2, 4 or 6 min (12 mW laser, Cobolt 06-mld). FOX solution (final concentrations: 0.25 mM ferrous ammonium sulfate, 25 mM methanolic $H_2SO_4$ and 0.1 mM XO in water/methanol 25:75) was added to the lipid suspensions and the absorbance was recorded from 450 nm to 700 nm. Control samples of lipids and Flipper (columns dubbed dark in Figure 1b) were left in the dark for additional 45 min after the 15 min incubation period and then the FOX solution was added. Statistical analysis was performed using GraphPad 10.2.2, *P* values less than 0.01 were considered statistically significant.

#### Singlet oxygen ($^1O_2$) detection

Lipid solutions of DOPC or DPPC at 50 µg/mL in $D_2O$ were incubated with Flipper (2.5 µM) for 15 min in the dark. ADMA (6 µM final concentration) was added and the solutions were illuminated for 30 seconds intervals with a 488 nm laser (12 mW, Cobolt 06-mld). The changes in the emission of ADMA were recorded ($\lambda_{ex}$ = 375 nm $\lambda_{obs}$ = 390 nm – 500 nm) using a Synergy H1 microplate reader (BioTek).

In $^1O_2$ quenching experiments, 10 mM $NaN_3$ was added to the solutions prior to light irradiation. Control dark and light experiments were carried out, where lipid solutions incubated with Flipper were kept in the dark for the duration of the experiments and lipids without Flipper were exposed to the same light dose, respectively.

### GUVs formation and imaging

GUVs were prepared following the PVA-assisted method with minor modifications.[1] Glass coverslips were cleaned by 15 min of sonication in acetone followed by 15 min in alkaline detergent (Hellmanex). Then, 40 µL of PVA solution (5 % w/w) in milliQ water was spread on the coverslip and dried for 30 min at 50 °C. A total volume of 10 µL of the corresponding lipid at 1 mg/mL doped with 0.02 % biotinylated and 0.1 % fluorescently labeled lipids (only in two color experiments) in $CHCl_3$ was spread on the PVA film and placed under vacuum for 1 h. The film was hydrated with 200 µL of 200 mM sucrose solution for 30 min at room temperature for DOPC and POPC, or 48 °C for DPPC and the ternary mixture POPC/DPPC/Chol. GUVs were collected using a cut pipette tip and diluted in 200 mM glucose solution, incubated with 1 µM Flipper (final concentration) for at least 15 min and imaged at room temperature in 18-well glass bottom dishes (Ibidi) previously coated with 50 µg/mL Neutravidin solution for 3 h at 37 °C or overnight at 4 °C.

### Confocal and FLIM imaging

Confocal and lifetime images were obtained using a TCS SP8 microscope (Leica Microsystems) equipped with an HC PL APO CS2 60x/1.40 oil objective, a pulsed white light laser (WLL) operating at 20 MHz repetition rate, 70 % master power and a hybrid detector (HyD) in photon-counting mode and 16-bit depth. Flipper was excited at 488 nm and the emission was collected in the 530-650 nm range, except when imaging vesicles co-labeled with far-red dyes, where detection was narrowed to 530-630 nm. Samples containing Atto647N or DiD were excited at 650 nm and the emission was detected between 670-750 nm. Lifetime measurements and analysis were performed using LAS X software. The lifetimes were calculated by fitting a bi-exponential decay and values from the longest decay were used.[2] Averaged lifetime images were obtained using the phasor plots. For phasor analysis, median or wavelet filters with a threshold of 5-8 were applied to better differentiate the photon clouds, which were then manually selected with linear or circular tools of radius 20-25.

Simultaneous FLIM and photo-oxidation experiments were performed at 488 nm by consecutive imaging scans with unidirectional exposure at 400 Hz and using 2x line accumulation, resulting in pixel dwell-times around 4-5.5 µs and a total time per frame of about 2.5 seconds. The power of the laser was typically set to 25 % of the master power, which resulted in 52 µW maximum power measured at the back of the objective using the bleaching point method provided from Leica, except when other powers were tested as indicated in the text (*see* Figure S2a). For dual-color imaging, sequential scan mode between lines was used.

### Cell culture

HeLa cells were grown in DMEM medium at 37 °C and 5 % $CO_2$. Transfection of Halo-tagged variants of GPI-AP and TfR was performed using the plasmids Str-KDEL_SBP-Halo-GPI and Str-KDEL_TfR-SBP-Halo, respectively, which carry the so-called retention using selective hooks (RUSH) system.[3] RUSH is based on the expression of streptavidin (Str) together with a targeting motif (in this case to the endoplasmic reticulum (ER) using the protein KDEL) and the reporter protein (GPI-AP or TfR) linked to a Str-binding peptide (SBP) motif. Once expressed, both proteins are retained in the ER until the addition of 40 µM D-biotin, which induces the displacement of SPB from Streptavidin and the release of the proteins. Cells were incubated at 37 °C for additional 60-90 min after biotin addition to ensure sufficient accumulation of the recombinant proteins at the plasma membrane and then incubated with the far-red Halo-JFX$_{650}$ (1 µM) for 15 min. Cells were washed three times with warm PBS to remove the unbound dye.

Preparation of GPMVs

GPMVs were produced and isolated from HeLa cells as described in Sezgin *et* al.[4] Cells were seeded on 35 mm glass bottom dishes (MatTek) and grown for 24 h to a confluency of around 70 %. To induce the formation of non-phase separated GPMVs, cells were washed twice with GPMV buffer (150 mM NaCl, 10 mM HEPES, 2 mM $CaCl_2$ at pH 7.4) and incubated with 25 mM PFA and 2 mM dithiothreitol for 2 h at 37 °C. Supernatants were collected and centrifuged for 90 seconds at 1000 g to remove detached cells. GPMVs were allowed to settle for 1 h and 20 µl from the bottom of the tube were diluted in 130 µL of GPMV buffer, incubated with 1 µM Flipper (final concentration) for at least 15 min and transferred to 18-well glass bottom chambers previously coated with 2 mg/ml BSA.

**Supplementary Figures.**

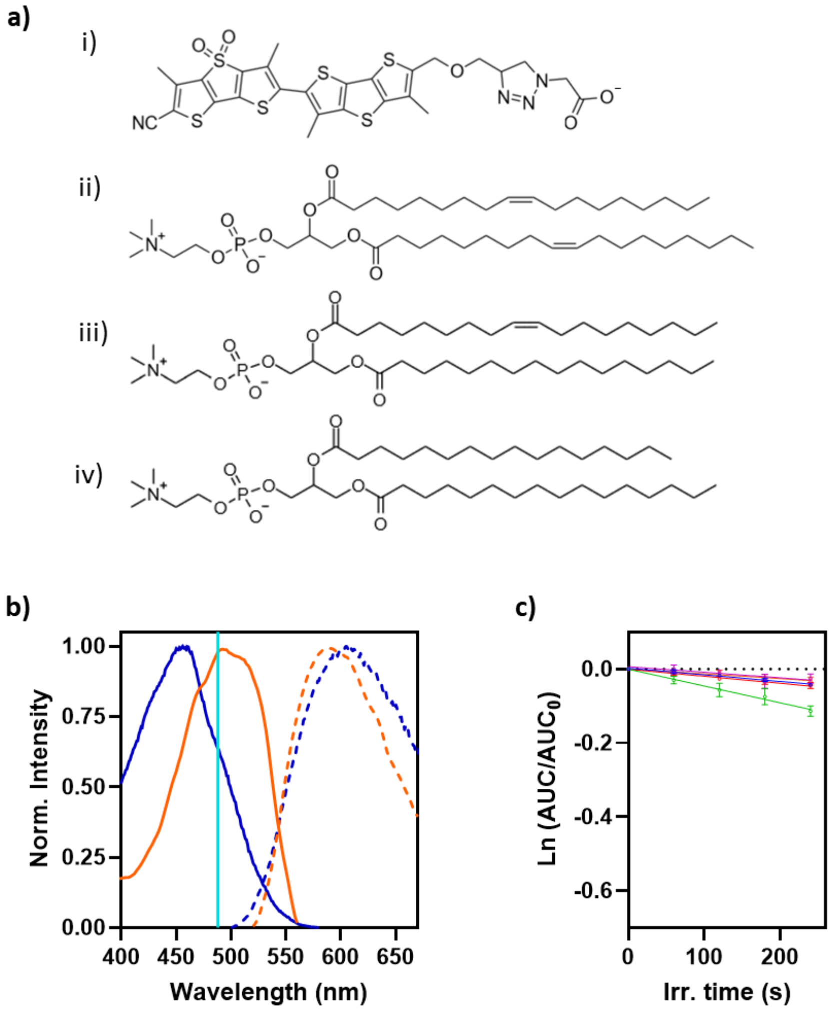


**Figure S1**. a) Molecular structures of (i) Flipper, (ii) DOPC, (iii) POPC and (iv) DPPC. b) Normalized fluorescence excitation (solid lines) and emission (dashed lines) spectra of Flipper in the presence of unsaturated DOPC (blue) and saturated DPPC (orange). Stronger planarization of the DTT scaffold of Flipper in DPPC membranes induces a red shift in the excitation spectra as compared to more fluid DOPC. The cyan vertical line at 488 nm indicates the wavelength of the laser used for Flipper excitation. c) ADMA bleaching rates in dark and light control $^1O_2$ detection experiments. Details: Irradiated samples of DOPC and DPPC in the absence of Flipper (blue and red, respectively), DOPC or DPPC incubated with Flipper and kept in the dark for the duration of the experiment (magenta and orange, respectively) and illumination of Flipper in the absence of lipids (green).

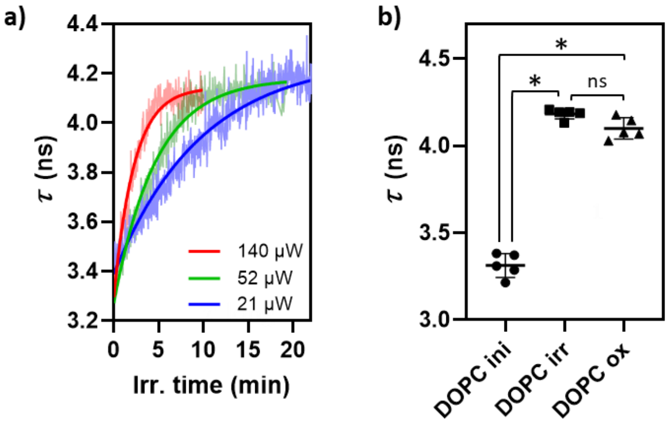


**Figure S2.** a) Time-course evolution of Flipper $\tau$ in DOPC GUVs under continuous FLIM imaging at different laser excitation powers. The transformations of the membranes that increase membrane tension occur in a light dose-dependent manner, reaching similar final $\tau$ values at the plateau. b) Flipper $\tau$ in DOPC GUVs before (DOPC ini) and after (DOPC irr) continuous FLIM imaging during at least 10 minutes, and in pre-hydroperoxidized DOPC (DOPC ox). DOPC irr values correspond to the average of the last 10 frames of the FLIM acquisition to exclude possible $\tau$ point fluctuations. Data shown are the mean ± SD of 5 replicates, **P* < 0.0001; ns: not significant, determined by unpaired one-way ANOVA .

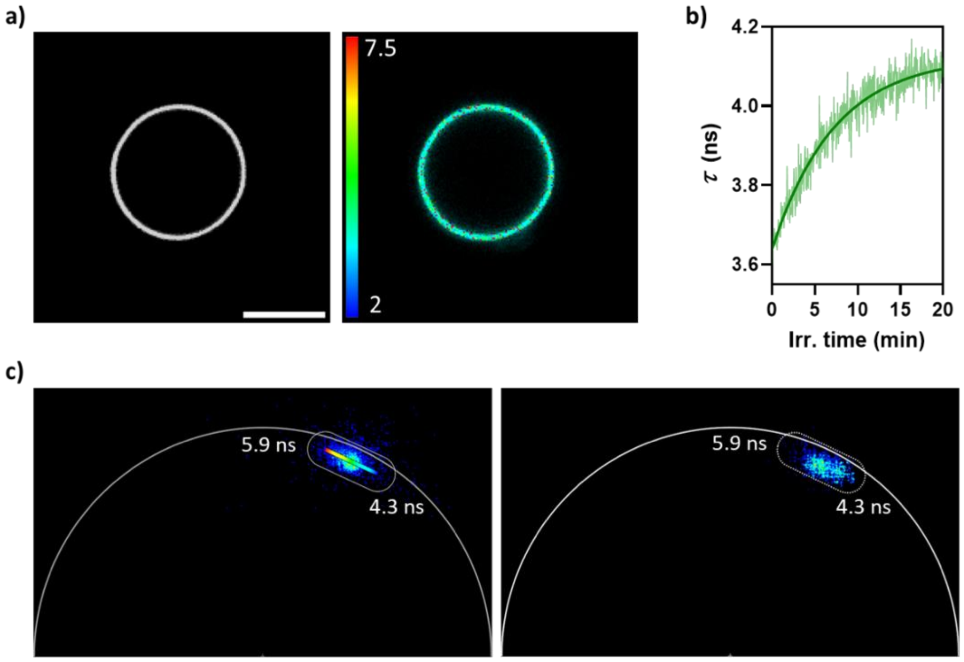


**Figure S3.** a) Intensity (left) and FLIM (right) image of a POPC GUV labeled with 1 µM Flipper showing a single phase and homogeneous $\tau$ = 3.64 ns. Scale bar is 10 µm. b) Continuous FLIM imaging at 488 nm (~52 µW) induces an increase in $\tau$ (reporting higher membrane tension), demonstrating the susceptibility of the lipid to photo-oxidation by Flipper upon blue light excitation. c) Phasor plots of Flipper in ternary POPC/DPPC/Chol (1:1:1) GUVs before (left) and after (right) phase separation induced by prolonged blue light Flipper excitation. The formation of small phase-separated domains creates a population of pixels at shorter averaged lifetimes. Phasor analysis using the linear tool was applied for better differentiation of the photon clouds and visualization of the segregated regions.

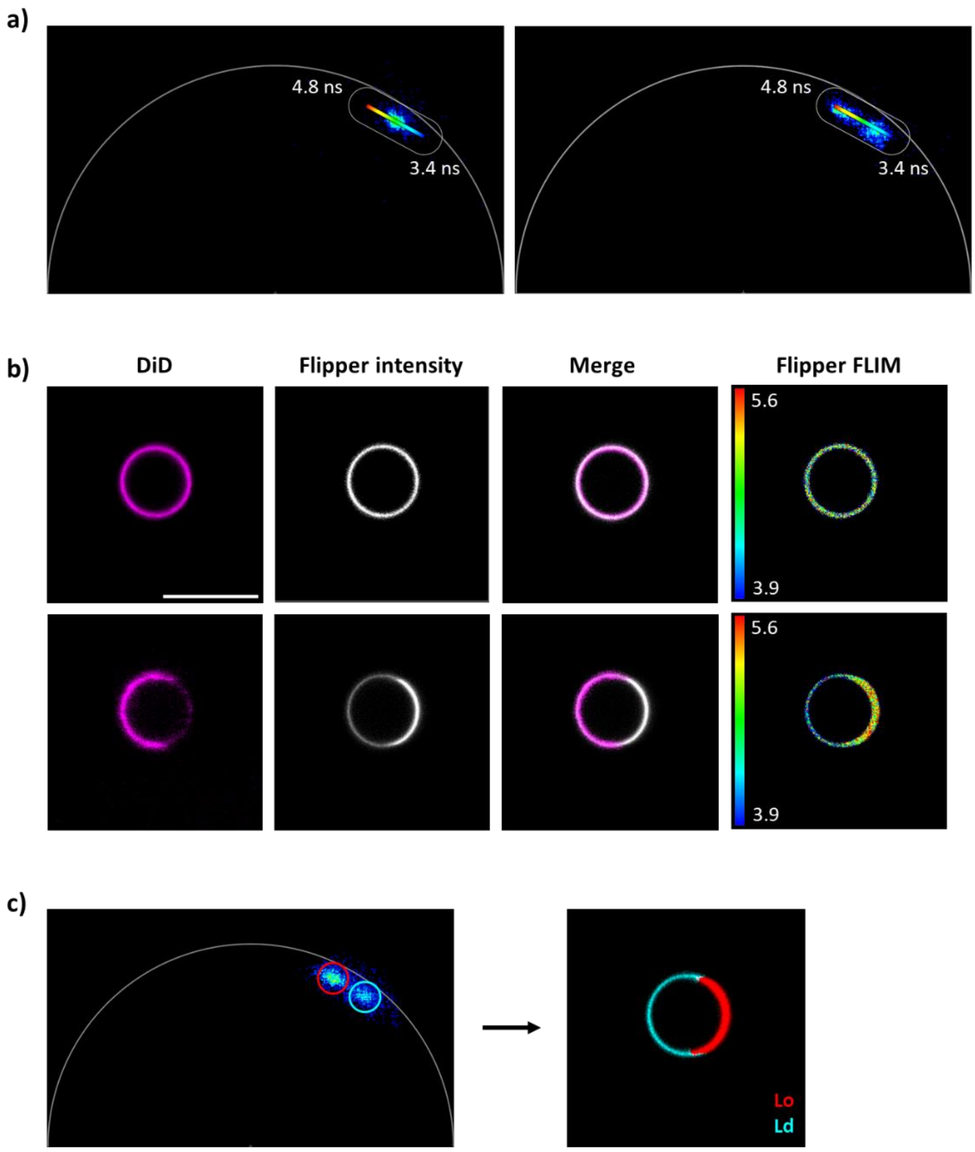


**Figure S4**. a) Phasor plots of Flipper in GPMVs before (left) and after (right) photo-induced phase separation. The transition from a single photon cloud centered around 4 ns to two well-differentiated populations enables the separation and visualization of the segregated domains using the linear tool from the phasor analysis. b) Dual-color images of a GPMV labeled with the Ld marker DiD and Flipper at initial times (upper row) and after phase separation triggered by prolonged blue light excitation (lower row), showing DiD accumulation in low-tension regions as revealed by weaker Flipper intensity and shorter $\tau$. c) Phasor analysis in photo-induced phase-separated GPMVs allows to define Lo and Ld domains using Flipper data.

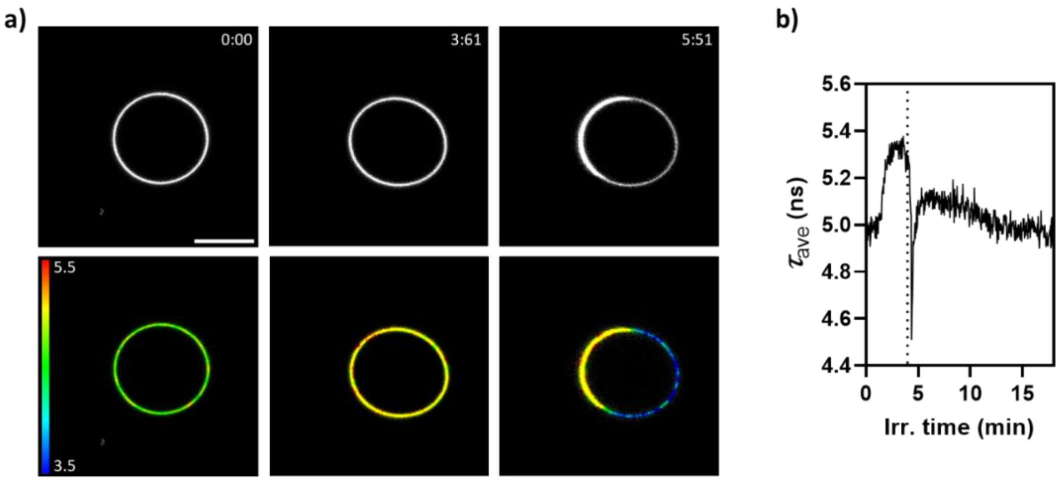


**Figure S5.** a) Selected frames over time of a GPMV labeled with Flipper under blue light excitation in consecutive FLIM imaging scans. An increase in $\tau$ (reporting higher membrane tension) is observed (second column) before the formation of segregated phases (third column). Irradiation times in min:sec are shown on the top right of the panels. Scale bar is 10 µm. b) Time-course of $\tau_{ave}$ collected from the whole GPMV reveals an increase in $\tau_{ave}$ prior to phase separation. The vertical dotted line corresponds to the first imaging frame where phase separation is observed.
